# Comparative effects of perfusate composition on rat brain histology following transcardial perfusion fixation

**DOI:** 10.64898/2026.09.16.752015

**Authors:** Autumn Beck, Andria Slaughter, Sarah Sedgewick, Macy Garrood, Katelyn Hedden, Andrew T. McKenzie

## Abstract

**Background:** Transcardial perfusion fixation is widely used to preserve rodent brains for histological and ultrastructural analysis, but protocols can vary in the use of pre-fixation washout, fixative formulation, and osmotic additives. Relatively few studies have directly compared how the chemical composition of the perfusate affects tissue preservation.

**Methods:** Male Sprague-Dawley rats underwent transcardial perfusion using a series of aldehyde-based fixative formulations that varied in fixative composition, use of a phosphate buffered saline washout, and the addition of mannitol or polyethylene glycol 35 kDa (PEG35). Perfusion outcomes were assessed using gross brain morphology, semi-quantitative grading of vascular blood clearance, quantitative detection of residual erythrocytes in whole-slide histological images, and light microscopic measures of cellular visualization and morphology. Perfusate osmolality was measured, and selected specimens were examined by electron microscopy.

**Results:** Substantial vascular blood clearance was frequently achieved with fixative-only perfusion, indicating that a pre-fixation washout was not required to achieve high levels of blood clearance under the conditions tested. Routine light microscopy-based measures of cellular visibility and morphology were broadly similar across treatment groups. Adding mannitol and PEG35 to the fixative solution both led to concentration-dependent gross tissue shrinkage, particularly with PEG35, without correspondingly large or consistent changes on light microscopy. A substantial degree of case-to-case variation in perfusion quality was also observed among animals undergoing nominally similar procedures.

**Conclusions:** Substantial variation in perfusate composition produced relatively modest differences in the light microscopy outcomes examined, despite pronounced effects of osmotic additives on gross brain morphology. There were some cases where perfusion with fixative alone led to near-complete blood vessel clearance, and the use of a preceding phosphate buffered saline washout was not associated with higher vascular clearance scores. The degree of case-to-case variability observed within treatment conditions suggests that procedural factors can play an important role on perfusion quality in addition to perfusate composition.

## Introduction

Transcardial perfusion fixation is the gold standard method for preserving whole rodent brains for histological and ultrastructural analysis [1,2]. It is widely relied upon in preclinical neuroscience for toxicology research, disease modeling, and therapeutic development, where high-quality tissue preservation is essential for accurate interpretation of experimental outcomes. Despite its importance, rodent perfusion fixation protocols vary considerably across laboratories, with differences in perfusion pressure and other variables [3,4]. Additionally, few studies have directly tested different possible perfusion fixation protocol parameters in a controlled setting in order to compare them. Many widely used protocols may be used primarily based on tradition rather than empirical evidence. As a result, there is a critical need to systematically evaluate how these different parameters affect preservation quality, which can be measured by the resulting tissue architecture as visualized via light microscopy.

One key question is to what extent a pre-fixation washout step, in which a non-fixative containing solution is perfused prior to perfusing a fixative-containing solution, is necessary for achieving high-quality perfusion. In a previous review of perfusion fixation in human brain banking, it was found that about half of studies employed a washout step, while many others reported satisfactory results without one [5]. Several additional sources have reported that the use of a washout solution is not necessary prior to perfusion fixation for achieving satisfactory tissue preservation [6–10]. Still other sources have reported that the use of a washout solution or a longer period of washout can in fact be deleterious, insofar as it delays the onset of fixation, which thereby decreases the tissue preservation quality [1,11–15]. On the other hand, many standard protocols for rodent perfusion fixation include a saline or PBS flush to clear blood cells and intravascular debris before fixative delivery [2,4]. The rationale for using a washout solution includes (a) preventing coagulation and associated clot-mediated obstruction of fixative flow and (b) preventing red blood cells (RBCs) from interfering in downstream assays. However, it is unclear if a washout solution is actually necessary to clear RBCs and achieve adequate fixative distribution throughout the brain, as the pressure-driven displacement of blood from the vasculature is much faster than the kinetics of aldehyde cross-linking [10]. To date, few studies have measured the degree of blood clearance when performing perfusion fixation with and without a washout step in rodent brains.

Another open question is to what extent the composition and concentration of the fixative perfused affects tissue preservation [16]. Two of the most commonly used aldehyde fixatives for transcardial perfusion are 4% paraformaldehyde (PFA) and 10% neutral buffered formalin (NBF), which contain approximately equivalent concentrations of formaldehyde (around 3.7% w/v) but differ in their additives, with NBF containing a small concentration of methanol as a stabilizer [17]. One study directly compared perfusion with 10% NBF and 4% PFA in a mouse model of experimental autoimmune encephalomyelitis, finding that the two fixatives produced similar histopathological outcomes, although the mice perfused with NBF showed a less severe burden of dark neuron artifact than those perfused with PFA [18]. However, there have not been a large number of direct comparisons between the use of NBF and PFA. Whether higher concentrations of fixative, such as 20% NBF, may improve perfusion quality is another question that has been suggested but not rigorously tested. Notably, one study in human brains reported that perfusion with 20% formalin led to more efficient fixation of deep brain structures compared to perfusion with 10% formalin [19]. The addition of glutaraldehyde to the fixative solution is standard practice for electron microscopy preparation, as glutaraldehyde cross-links proteins more rapidly than formaldehyde, which is thought to improve the fidelity of ultrastructural preservation [20,5]. Although several studies have used glutaraldehyde in protocols that have achieved high quality ultrastructural preservation [5,21], the effects of adding glutaraldehyde on perfusion and histology quality are still not fully established.

Finally, the extent to which the addition of osmotically active agents to the perfusate can affect fixation quality is relatively unexplored. This has been proposed as a method for ameliorating perfusion impairment in ischemic brains, but whether it also leads to effects on structural preservation is important for evaluating its utility for that purpose [22]. Previous research has reported that hypertonic aldehyde perfusates led to cellular shrinkage and expanded extracellular space, while isotonic and hypotonic solutions caused brain swelling and poor ultrastructural preservation [23]. Mannitol, a membrane-impermeant sugar alcohol commonly used as an osmotic agent in clinical neurology, is of particular interest in this context. One study developed a multi-step transcardial perfusion protocol in which mannitol was used at varying concentrations to first open the blood-brain barrier (15%), then to restore extracellular space (4.5%), and finally maintain the extracellular space during aldehyde fixation (4%) [24]. In the context of human brain banking, mannitol has been used as an additive to the washout or fixative solution, in an attempt to mitigate tissue edema [19,25,26]. However, the effects of adding mannitol at various concentrations to a standard formaldehyde-based perfusate on perfusion quality have not been systematically evaluated, and there is a concern that higher concentrations could induce excessive tissue dehydration. Colloids such as high molecular weight (MW) polyethylene glycol (PEG) are also of potential interest, as these have been used to improve perfusion quality in organ preservation and resuscitation contexts [22,27]. One study reported that the addition of a colloid (2% polyvinylpyrrolidone, MW 40,000) during perfusion fixation slightly improved ultrastructural preservation in brain regions with a blood-brain barrier, suggesting that colloids may help to maintain vascular integrity [28]. However, whether high-MW PEG also affects neurohistologic preservation quality has, to the best of our knowledge, not yet been thoroughly investigated.

These methodological questions in brain preservation are rarely addressed directly, because investigators are often more interested in downstream aspects of the research. In this study, we compared several methodological parameters of transcardial perfusion for brain preservation in rats. The variables included a pre-fixation washout step, fixative composition, fixative concentration, and the addition of osmotically active additives to the fixative solution.

Preservation quality was assessed through examination of the brain’s gross morphological appearance, histological assessment of vascular blood clearance, cellular morphology on light microscopy, and targeted ultrastructural imaging via electron microscopy. Our goal was to identify how the protocol modifications we tested influence measurable aspects of tissue preservation.

## Methods

### Laboratory animals

Male Sprague-Dawley rats (226-300 g; Charles River Laboratories, Wilmington, MA, USA) were purchased and housed with *ad libitum* access to food and water until the time of the experiment. The protocol was approved by the Sparks Brain Preservation Institutional Animal Care and Use Committee (Protocol #: IACUC-001; approved August 7, 2025). The procedures were conducted in accordance with the approved protocol and with accepted principles for the ethical care and use of laboratory animals. All terminal procedures were performed under deep isoflurane anesthesia in order to minimize animal suffering.

### Transcardial perfusion

Animals were anesthetized using 4% isoflurane delivered by inhalation. Adequate depth of anesthesia was confirmed by the absence of withdrawal to toe pinch before beginning the surgical procedure. Transcardial perfusion was performed as a terminal procedure under deep isoflurane anesthesia, and animals did not regain consciousness.

Transcardial perfusion was performed utilizing a peristaltic pump with an 18 gauge needle primed with the perfusate appropriate for the experimental condition (**Table 1**). The perfusate was placed in a 500 ml beaker and the tubing was secured inside of it. To begin, rats were positioned supine on a large wax dissection tray. From here, standard surgical scissors were used to cut a u-shaped midline incision through the abdominal wall just below the rib cage, and reflected up, so that a window was available to access the diaphragm from below for cutting. Then these scissors were used to cut through the diaphragm, after which we further cut through the rib cage on both sides. The thoracic cage was then folded up above the shoulder and pinned to the tray to maintain it in place and fully reveal the heart and thoracic cavity. This procedure took between 15-30 seconds.

**Table 1.**
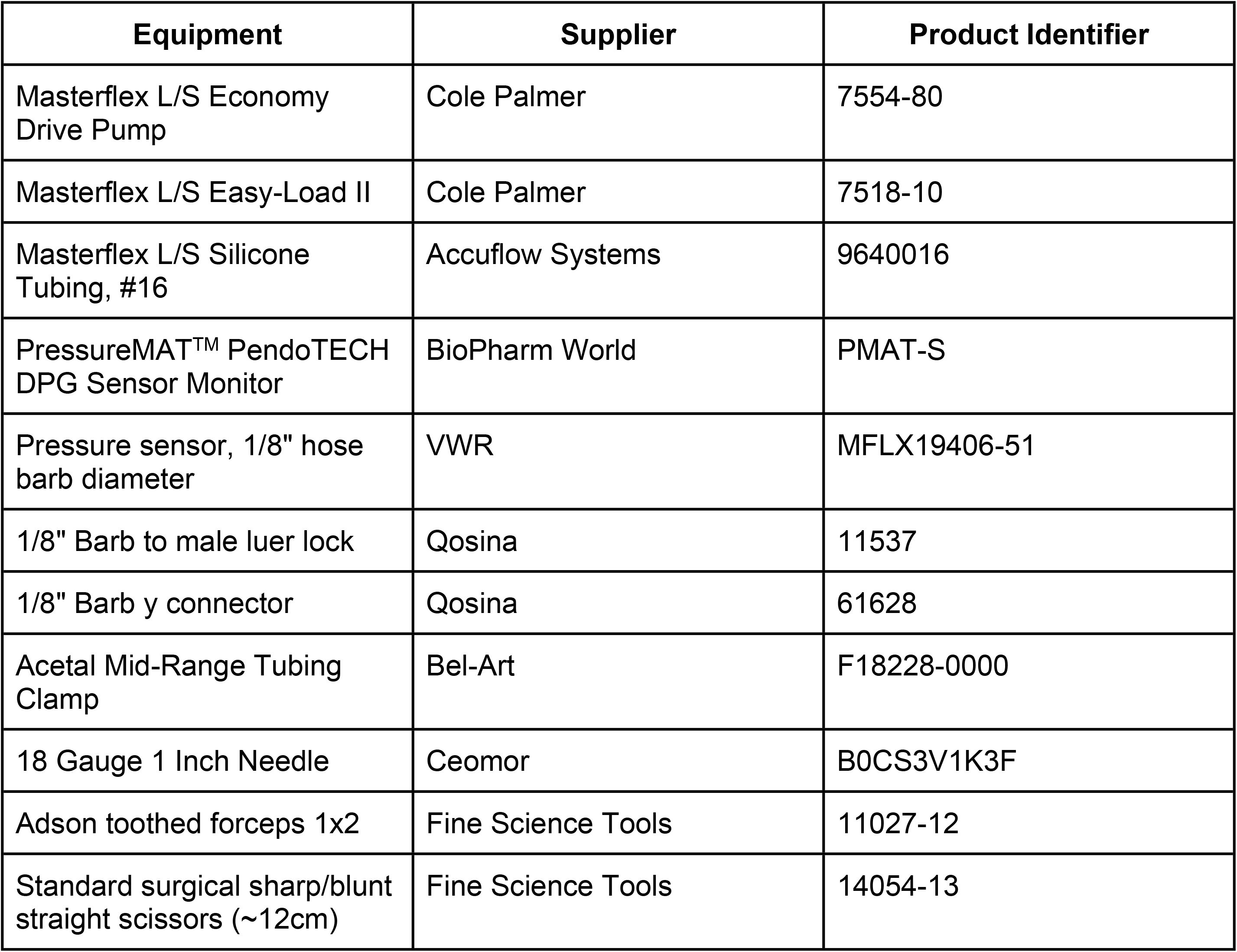
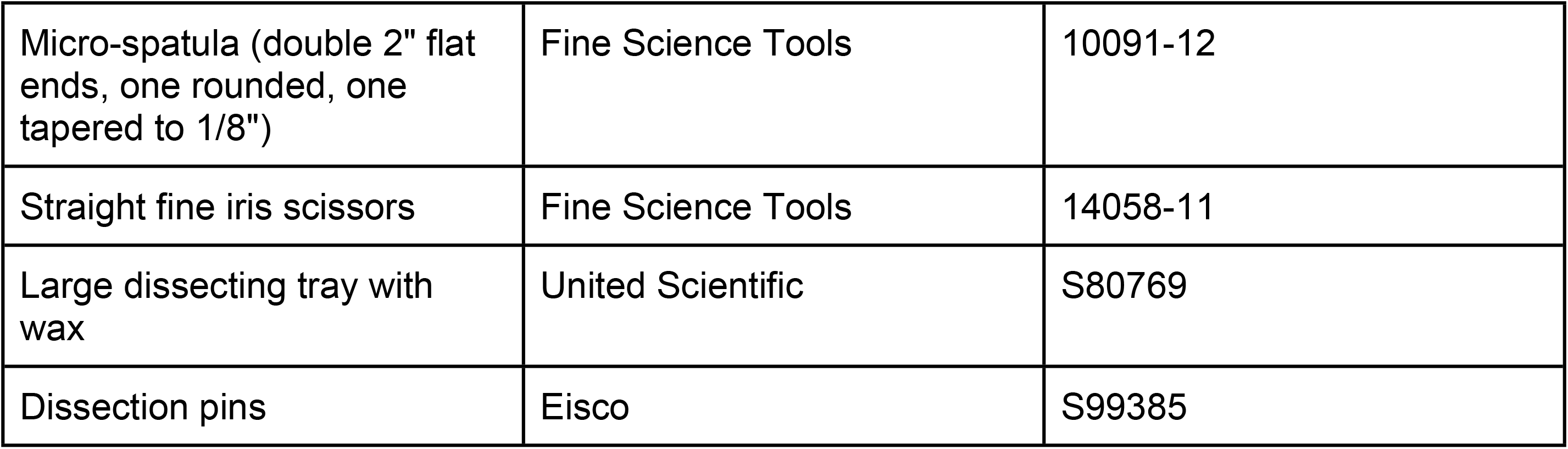
Supplies used for perfusion fixation and brain extraction.

An 18 gauge needle was then inserted into the inferior aspect of the left ventricle, directed towards the aorta. The right atrium of the heart was immediately clipped with iris scissors to allow for blood drainage and eventual perfusate drainage. The perfusate flow was then started at an initial flow rate of approximately 8 ml/min (corresponding to a speed control dial setting of 0.5 on the perfusion pump) for approximately 1-2 minutes. The speed was then increased by increments of 0.5 until it was assessed that further increases in speed would lead to a high risk of backflow of perfusate through the cannulation site because of excessive local pressure. With our perfusion circuit, a speed control dial setting of 1 corresponds to an approximate flow rate of 19 ml/min, 1.5 to 28 ml/min, and 2 to 46 ml/min (which was the maximum speed used in these experiments). Throughout the process, the pressure was monitored with a PressureMAT sensor and we recorded the pressure whenever the flow rate was changed. The return flow from the right atrium became clear after a few minutes of perfusion in every case. The perfusion was continued until approximately 175-375 ml (more commonly 200-325 ml) of fixative solution was perfused. After perfusion, the inner lobes of the liver were generally noted to be pale and the whole body was generally stiff.

For experiments incorporating a washout step, the perfusion tubing was connected via a Y-connector to two separate 500 mL beakers containing washout solution and fixative (**Table 2**). The fixative line was primed first and then clamped. The washout line was subsequently primed, displacing the residual fixative from the shared tubing. The perfusion was initiated with the washout line unclamped and the fixative line clamped. At the completion of the washout (which was either approximately 50-60 ml or approximately 150 ml), the perfusion was switched to fixative by first unclamping the fixative line and then clamping the washout line.

**Table 2.** Chemical reagents used.

| Chemical | Supplier | Product Identifier |
| --- | --- | --- |
| 10% neutral buffered formalin (NBF) | Azer Scientific | NBF55G |
| 20% neutral buffered formalin (NBF) | Fisher Scientific | STL286205 |
| Phosphate buffered saline (PBS) Tablets | Fisher Scientific | BP2944-100 |
| Glutaraldehyde, 50% aqueous solution | Fisher Scientific | A10500 |
| Mannitol Powder USP Grade 12.5 kg | Lab Alley | MANPU-12.5KG |
| Polyethylene glycol, 35,000 average MW (PEG35) | Sigma-Aldrich | 81310 |

### Preparation of perfusates

Perfusates were generally prepared immediately prior to use. In some cases, they were prepared within 24 hours of use and stored temporarily prior to perfusion. In this case, they were generally stored at room temperature, with the exception of solutions containing glutaraldehyde, which were stored at refrigerator temperature prior to use. For experiments using glutaraldehyde supplementation, we used a final glutaraldehyde concentration of 1%, made from a 50% aqueous glutaraldehyde stock solution, which was added to 20% NBF. Mannitol and PEG35 were mixed with NBF with a magnetic stirrer for approximately 5-15 minutes or until visually homogenous, using some degree of heating (to approximately 50°C) to accelerate the process of mixing.

### Osmolality testing

The perfusate osmolality was measured by freezing point depression using an OsmoTECH XT Single-Sample Micro-Osmometer (Advanced Instruments, Norwood, MA, USA; **Table 3**). Measurements were performed in quintuplicate.

**Table 3.**
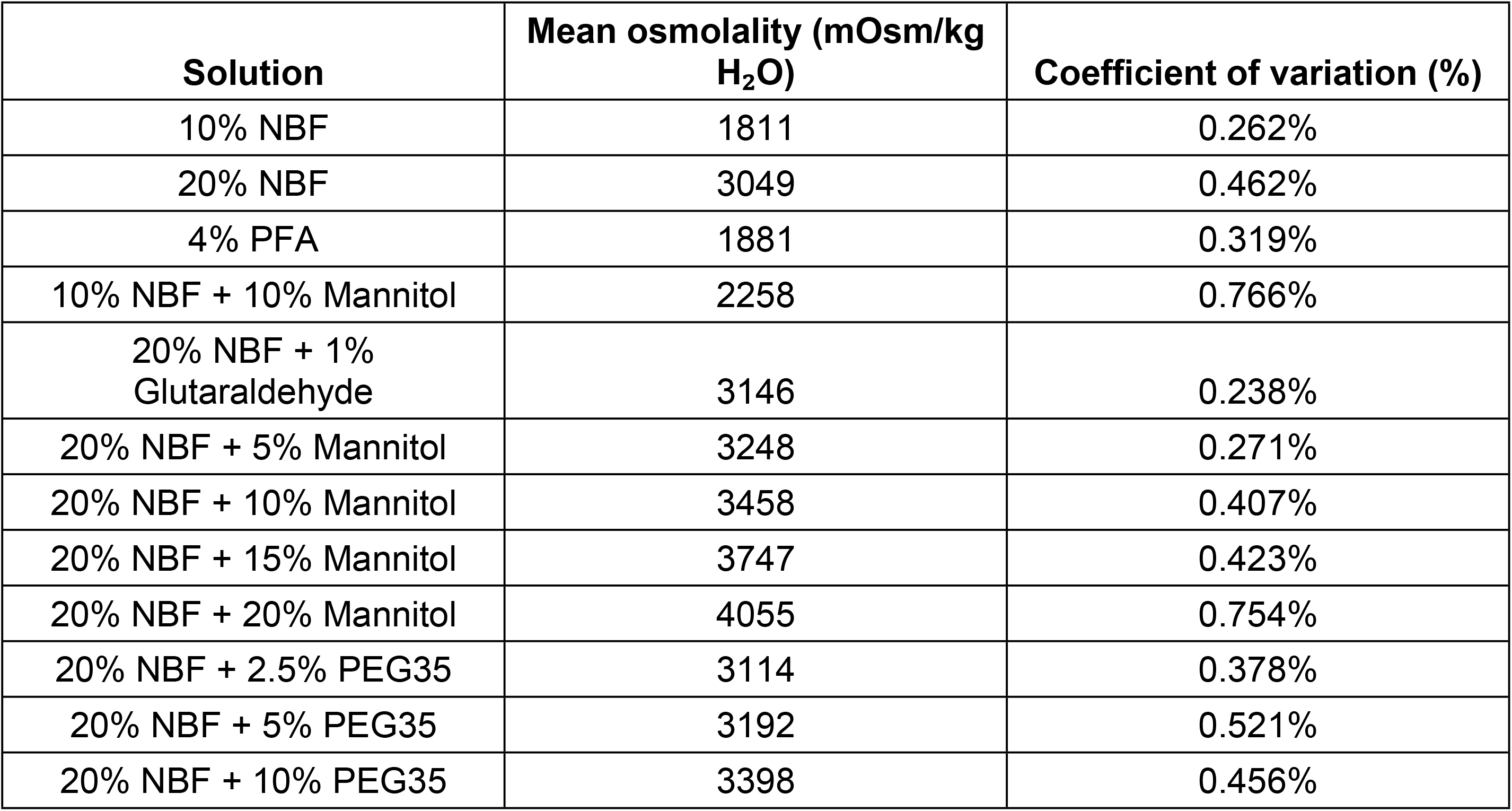
Measured osmolality of perfusate formulations determined by freezing point depression. Values represent the mean of five measurements unless otherwise indicated.

| Solution | Mean osmolality (mOsm/kg H<sub>2</sub>O) | Coefficient of variation (%) |
| --- | --- | --- |
| 10% NBF | 1811 | 0.262% |
| 20% NBF | 3049 | 0.462% |
| 4% PFA | 1881 | 0.319% |
| 10% NBF + 10% Mannitol | 2258 | 0.766% |
| 20% NBF + 1% Glutaraldehyde | 3146 | 0.238% |
| 20% NBF + 5% Mannitol | 3248 | 0.271% |
| 20% NBF + 10% Mannitol | 3458 | 0.407% |
| 20% NBF + 15% Mannitol | 3747 | 0.423% |
| 20% NBF + 20% Mannitol | 4055 | 0.754% |
| 20% NBF + 2.5% PEG35 | 3114 | 0.378% |
| 20% NBF + 5% PEG35 | 3192 | 0.521% |
| 20% NBF + 10% PEG35 | 3398 | 0.456% |

### Experimental design

We evaluated several variables that may influence the quality of transcardial perfusion fixation, including the use of a pre-fixation washout step, fixative composition, fixative concentration, and the addition of osmotically active agents to the perfusate (**Table 4**). Notably, these groups are a series of related comparisons rather than a fully crossed factorial design.

**Table 4.** Assignment of rats to experimental groups. Rat identification numbers are listed for each perfusion or immersion fixation condition.

| Perfusate group | Rat numbers |
| --- | --- |
| 10% NBF | 1, 57, 58 |
| 20% NBF | 6, 7, 8, 12, 17, 22, 23, 30, 31, 32 |
| 10% NBF and 10% Mannitol | 2, 3, 4 |
| 20% NBF + 1% Glutaraldehyde | 9, 10, 11 |
| 20% NBF + 5% Mannitol | 13, 18, 24, 25, 33 |
| 20% NBF + 10% Mannitol | 14, 19, 26, 27, 34 |
| 20% NBF + 15% Mannitol | 15, 20, 28, 29, 35 |
| 20% NBF + 20% Mannitol | 16, 21, 36 |
| 20% NBF + 2.5% PEG35 | 37, 38, 42, 43 |
| 20% NBF + 5% PEG35 | 39, 40, 44, 45 |
| 20% NBF + 10% PEG35 | 41, 46, 47, 48 |
| 4% PFA | 49, 50, 53 |
| Washout with PBS (50-60 ml), then 20% NBF | 54, 55, 56 |
| Washout with PBS (150 ml), then 20% NBF | 59, 60, 61 |
| Immersion fixation in 10% NBF | 5 |
| Immersion fixation in 20% NBF | 51, 52 |

The animals were not randomly assigned to experimental conditions. Instead, treatment conditions were selected pragmatically over the course of the study, with some interleaving between experimental groups. Sample sizes were determined during the course of this exploratory study, generally aiming for at least three animals per perfusion condition. No *a priori* power calculation was performed. A total of 61 rats were used in the study.

The individual rat was considered the experimental unit for comparative analysis. Because this was an exploratory methodological study, the measured outcomes were treated as complementary exploratory measures of perfusion and preservation quality rather than as a single prespecified primary endpoint.

No *a priori* exclusion criteria were specified. Exclusions made during analysis are described below. Rats 1-4 and 10 were not included for histology analyses because these brain samples had already been sectioned during pilot work, so there was not enough tissue available. Rat 56 was excluded from the vascular clearance analysis because the perfusion in this case totally failed, such that the specimen was not considered an example of adequate delivery of the assigned washout and fixation protocol. This exclusion decision was based on the observed perfusion failure, as opposed to the subsequent vascular clearance measurement.

### Brain extraction

Cephalon isolation was completed in 30 seconds or less using standard surgical scissors by cutting the cervical vertebrae from the posterior angle. The cephalon of the rat was then pinned to a wax tray and the soft tissue was removed to expose the superior and lateral surfaces of the skull. If the atlas or axis cervical vertebrae were still attached at the base of the skull, they were removed via Adson toothed forceps, exposing the foramen magnum and occipital bone. Careful insertion of fine scissors mediolaterally into the foramen magnum was then done to snip into the occipital bone, giving break-away points for the skull as it is fragmented away from the brain into pieces with Adson toothed forceps. Once at the superior portion of the skull, fine scissors were used to cut a central line along the sagittal suture as well as the interfrontal suture. Adson forceps were used to fragment large pieces of the temporal, parietal, and frontal bone away from the brain. Once exposed, a micro-spatula was used to gently break away cranial nerves and arteries, thus finalizing the isolation of the brain. Following extraction, digital photographs of the brain were taken from multiple angles.

These photographs were taken prior to immersion fixation, to prevent this process from confounding any color changes in the brain. Rat 61 was excluded from the gross image analysis because post-extraction photographs were not available. After the photographs were taken, the brains were completely immersed in the same fluid as was used for perfusion, which was then stored at 4°C until use [29]. The features used for estimating the extent of perfusion on the images were (a) the clearance of blood from surface blood vessels, (b) the apparent stiffness of tissue based on its texture, and (c) any color changes of tissue, from pink in the case of no perfusion to pale or in the case of well-perfused tissue. The same gross examination images were graded independently by two raters.

### Histology

Brain samples were manually blocked in the coronal plane with the goal of obtaining comparable transverse sections across specimens. Because blocking was performed manually, the precise anatomical level varied somewhat between brains. The blocks were then placed into cassettes for processing and embedded in paraffin. Paraffin-embedded brain sections 6 µm thick were baked, deparaffinized, and stained for Luxol Fast Blue, Hematoxylin, and Eosin (LH&E). Digital images of the stained sections were captured at 40X as whole slide images (WSIs) using the Aperio GT450 high-resolution scanner (Leica Biosystems).

### Vascular clearance

To analyze the histology data semi-quantitatively, two graders independently assessed the extent of vessel clearance across each of the WSIs on a 0-3 scale, with 0 indicating <5% clearance of blood vessels, 1 indicating 5-50%, 2 indicating 50-95%, and 3 indicating >95%, as we previously performed [26]. Vessel clearance refers to the absence of intravascular material from both small and large vessels, as in some cases there was clearance from the large vessels but not the small vessels. These grades were then compared via quadratic-weighted Cohen’s kappa, and the grades from the two reviewers were averaged for downstream analysis.

### Red blood cell quantitative assessment

Red blood cells (RBCs) were quantified across WSIs using a pixel classifier trained in QuPath (v. 0.7.0). The classifier was a random trees pixel classifier implemented through OpenCV and used the red, green, and blue image channels smoothed with a Gaussian filter (sigma = 1.0) as input features. Classification was performed at a resolution of 2.1 µm per pixel using 512 × 512 pixel tiles. Pixels were classified as either RBC-positive or background.

Classification accuracy was improved by annotating RBC-positive regions primarily within the central portion of each RBC while excluding the darker peripheral edges of the cells. Training annotations were initially concentrated on the WSI from Rat 6. However, classifier training was performed progressively, and additional annotations were subsequently added from each WSI to improve classifier performance across slides.

The trained classifier was applied to all available WSIs within standardized bounding boxes of approximately 500,000 µm². Bounding boxes were centered within the tissue to avoid tissue edges and to maintain consistent analysis areas across WSIs. Two anatomical regions were evaluated, which were called Parent Boxes 1 and 2. Parent Box 1 was positioned in the central region of the brain and contained mixed gray and white matter, whereas Parent Box 2 was positioned closer to the tissue edge within the cerebral cortex and primarily contained gray matter.

RBC abundance was quantified within each bounding box using multiple metrics that were measured using the QuPath script editor. First, the RBC density was defined as the percentage of pixels within each parent box classified as RBC-positive. Second, the number of discrete RBC-positive objects within each parent box was also recorded, also known as the annotation count. Third, the cumulative area occupied by RBC-positive detections within each parent box was calculated. This value was expressed as a percentage of the total parent box area and was defined as the RBC area fraction.

Following a manual review of classifier performance across all WSIs, Rats 51, 54, 55, and 57 were excluded from subsequent classifier-based analyses, because the RBC detection in these cases was found to be unreliable. Specifically, these slides had a pronounced blue hue within the RBCs, likely resulting from variation or artifacts introduced during the staining process, which interfered with accurate RBC classification.

### Cellular morphology assessment

For each WSI, a rectangular region of interest (ROI) with an area of 630,000 μm² was placed within the cortical layers overlying the white matter. The white matter was identified by its Luxol fast blue staining. Using QuPath’s positive cell detection function, cellular profiles were detected and recorded as objects. Under setup parameters, the detection image was set to optical density sum with requested pixel size = 0.5 μm. The nucleus parameters remained as follows: Background radius = 8 μm, Median filter radius = 0 μm, Sigma = 1.5 μm, Minimum area = 10 μm², Maximum area = 400 μm². The intensity parameters (threshold and max background intensity) were adjusted in accordance to the intensity of staining in each image. The range of intensity of the threshold was from 0.02-0.15 while max background intensity ranged from 1 to 2. The split by shape option was selected and the exclude DAB option was not selected. The cell parameters were set to exclude cell nucleus visualizations, with cell expansion set to 0 μm. Under general parameters, Smooth boundaries and Make measurements were selected. Under intensity threshold parameters, the score compartment was selected as Nucleus: DAB OD mean with Threshold 1+ = 0.1, Threshold 2+ = 0.4, Threshold 3+ = 0.6, and the Single threshold option selected. Objects that were identified as background artifacts, noise, or blood cells were manually reviewed and removed prior to export of data. Polygon vertices describing the boundaries of detected objects were exported from QuPath for subsequent morphometric analysis.

Two reviewers independently graded each ROI based on two measures. First, the ability of the QuPath model to accurately outline the shape of each cellular profile, on a 0-3 scale. Errors in this could be due to not detecting cells or inaccurately outlining the shape of the cell. Second, the visibility of the cellular profiles compared to the background, which was also graded on a 0-3 scale. This was not based on QuPath’s model but just based on the WSI tissue quality, including the staining quality. For each of the two measures, the two independent grades were averaged for downstream analysis.

Polygon vertex data exported from QuPath were analyzed in Python (v. 3.14) to calculate morphometric features and classify each detected object into one of four categories, evaluated hierarchically in the following order: (a) Partial detection, (b) Other, (c) Elongated, and (d) Round. The first matching criterion determined the assigned category. Objects with solidity <0.75 were classified as partial detections (often, these had a crescent shape). Of the remaining objects, those with compactness >1.6 were classified as other, and those with an aspect ratio >1.5 were classified as elongated (often, these had an oval shape). All remaining objects were classified as round.

### Electron microscopy

Tissue was post-fixed and processed using an adapted NCMIR heavy-metal staining protocol [30]. This process included sequential treatments with tannic acid, reduced osmium, thiocarbohydrazide, osmium, and uranyl acetate at room temperature, followed by lead aspartate staining at 60°C. Samples were dehydrated through graded ethanol and acetonitrile, infiltrated Embed 812 epoxy resin (EMS), and polymerized for 72 hours at 60°C. Ultrathin sections (70-110 nm) were cut using a Leica UC7 ultramicrotome and collected on glue-prepared copper slot grids. Electron microscopy images were acquired using a Zeiss Crossbeam 350 in scanning transmission electron microscopy (STEM) mode.

### Grading methodology and statistical analysis

Reviewers were not formally blinded to treatment allocation, although treatment labels were not displayed during grading. Statistical analyses were performed primarily in Google Sheets. Inter-rater agreement for ordinal grading scales was assessed using the quadratic-weighted Cohen’s kappa. Given the small group sizes and the ordinal nature of several outcomes, nonparametric statistical tests were used and no assumption of normality was made. Comparisons between two independent groups were performed using the Mann-Whitney U test. Comparisons across multiple treatment groups were performed using the Kruskal-Wallis test, followed, where appropriate, by Dunn’s post hoc pairwise comparisons. Associations between continuous or ordinal variables were assessed using Spearman’s rank correlations. All statistical tests were two-sided. To account for multiple hypothesis testing within sets of related exploratory analyses, the Benjamini**-**Hochberg procedure was used to control the false discovery rate at 0.05. For analyses subjected to multiple hypothesis testing correction, statistical significance was determined using the Benjamini-Hochberg adjusted results, while raw p-values are reported where indicated. Dunn’s post hoc pairwise comparisons were likewise adjusted for multiple comparisons using the Benjamini-Hochberg procedure. A two-sided p-value of <0.05 was considered statistically significant.

## Results

### Gross examination

Gross photographs obtained immediately after brain extraction and before immersion fixation were independently evaluated by two reviewers. The apparent distribution of perfusate was graded on a 0-3 scale based on the degree of residual red or pink tissue coloration, visible blood within superficial vessels, focal blood pooling, and the uniformity of pallor across the brain surface (**Figure 1**). The inter-rater agreement for these grades was high (quadratic-weighted Cohen’s κ = 0.84, n = 60). Most perfused brains showed gross evidence of successful perfusion, including widespread pallor of the brain surface and reduction of visible blood within superficial vessels. However, several of the brains showed some degree of mottling, i.e. patchy areas of residual pink coloration across an otherwise relatively pale brain surface (e.g. **Figure 1C**). We also noted a tendency for some of the brains to have incomplete perfusion in the lateral surfaces of the cerebral hemispheres, as indicated by a pink-red color in these regions, for unclear reasons (**Figure 2**).

**Figure 1.**
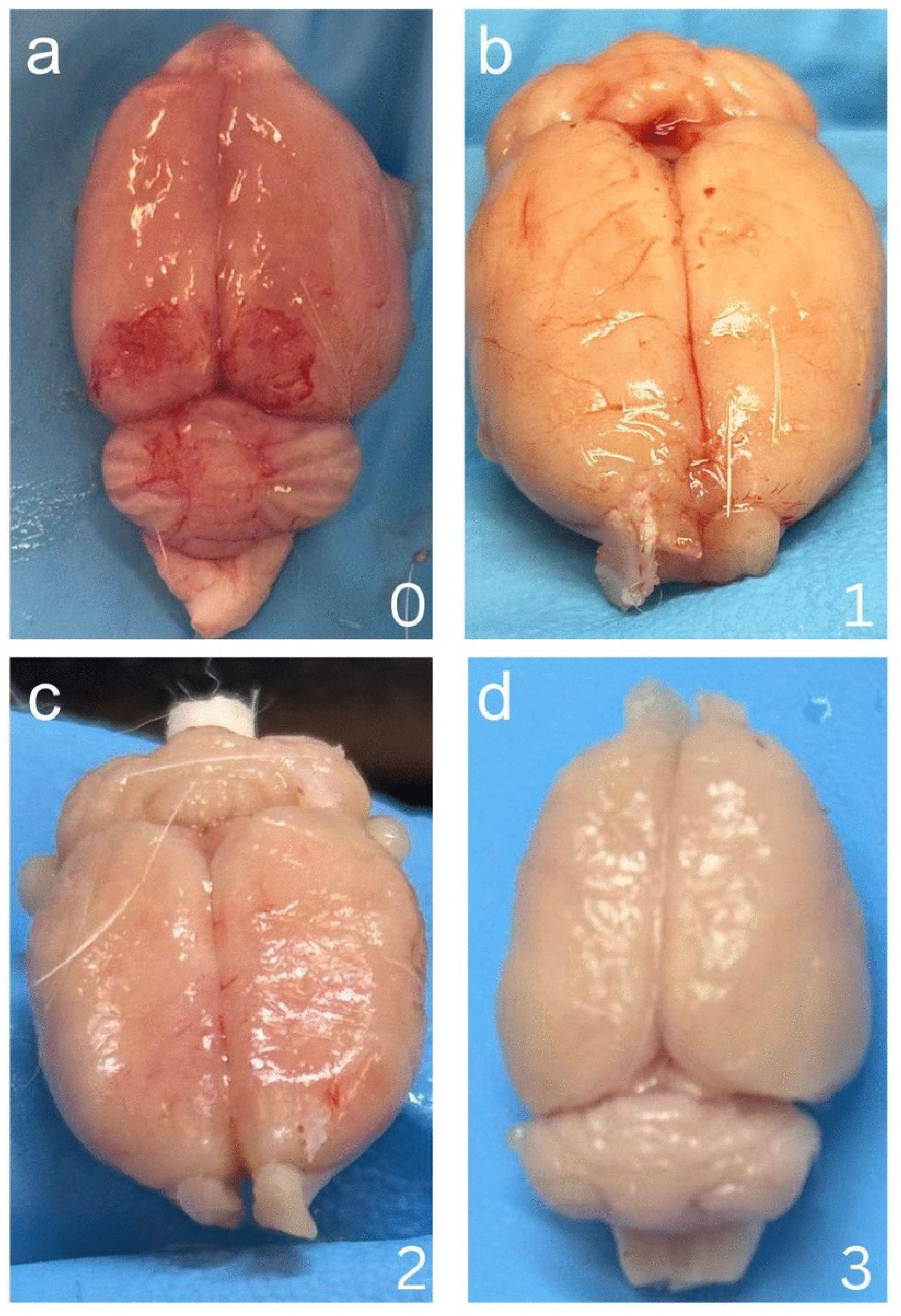
Representative macrostructure quality scoring of perfused rat brains. Brains were imaged immediately after extraction and prior to immersion fixation. They were assigned a perfusion quality score of 0-3 based on their gross appearance: 0, dark red or pink tissue with a substantial degree of visible blood pooling (**a**); 1, residual tissue redness and blood visible within the superficial vasculature (**b**); 2, predominantly pale-pink tissue that may have a small amount of visible residual blood (**c**); and 3, uniformly pale or white tissue with little or no visible residual blood (**d**). The representative image from the 0 score is from Rat 5 (which was immersion fixed), 1 from Rat 45, 2 from Rat 44, and 3 from Rat 7.

**Figure 2.**
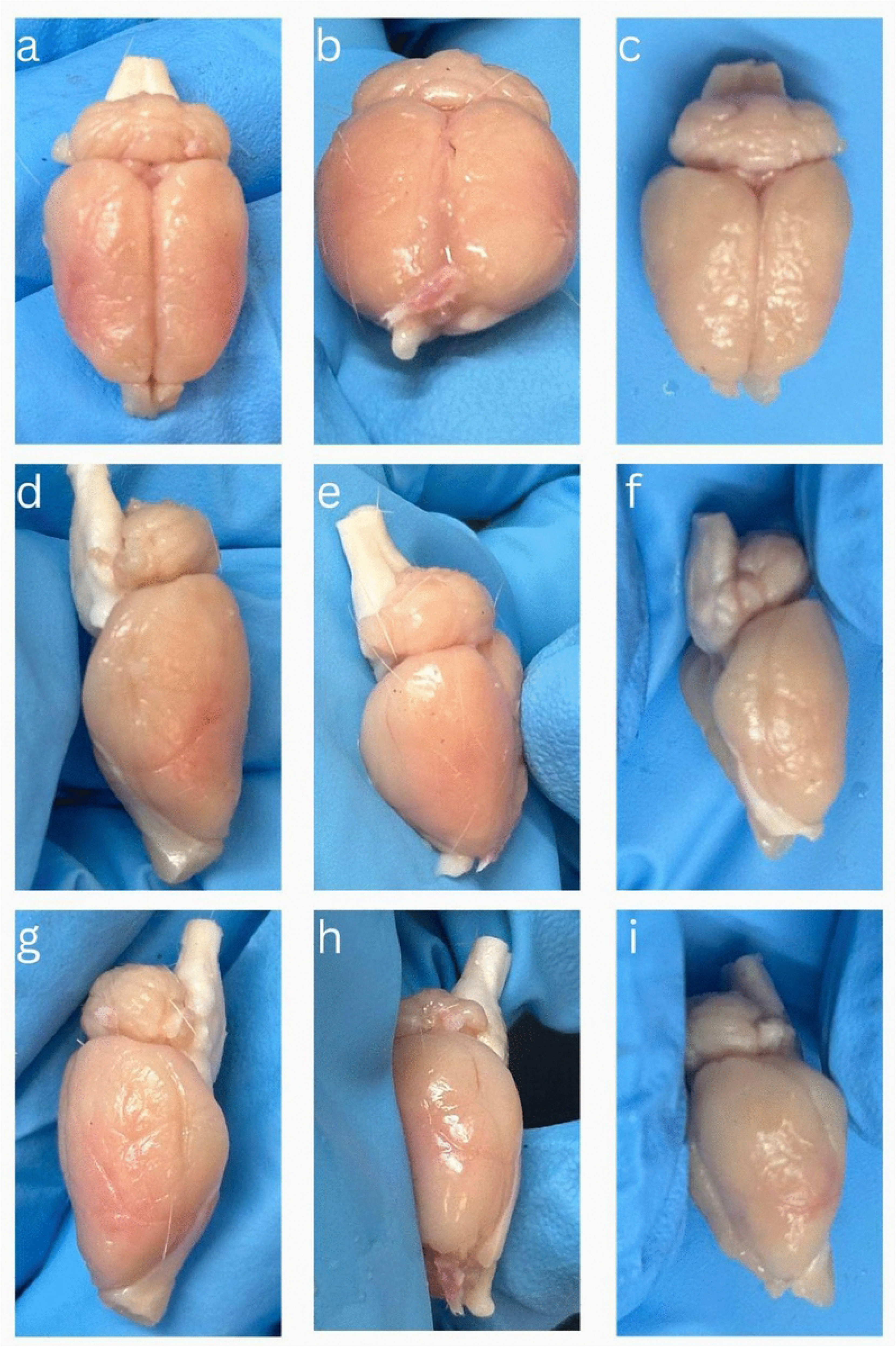
Representative examples of residual lateral blood following brain perfusion. Each column shows multiple views of the brain from a single rat: Rat 7 (**a**, **d**, **g**), Rat 59 (**b**, **e**, **h**), and Rat 12 (**c**, **f**, **i**). Brains were photographed immediately after extraction and prior to immersion fixation. In each animal, residual pink-red coloration is visible along portions of the lateral cerebral surface, despite otherwise relatively pale gross appearance, indicating incomplete perfusion of these areas.

Qualitatively, we found that there was not a substantial difference in the degree of blood clearance in the groups that did and did not include a washout step. We had numerous brains that were directly perfused with fixative solution and had apparent full clearance of blood on macroscopic examination, indicating that the use of washout solution is not necessary to clear blood from the vascular system in this context, as assessed by gross examination.

We also found qualitatively that the addition of mannitol or PEG35 produced clearly apparent changes in gross brain morphology. Increasing concentrations of mannitol in 20% NBF were associated with a progressively greater tissue contraction and a more dehydrated appearance (**Figure 3**). These changes were most apparent at the higher mannitol concentrations, particularly 15% and 20%, while brains perfused with lower concentrations showed a less pronounced gross deformation. In brains perfused with higher concentrations of mannitol, free fluid was also observed within the cranial cavity surrounding the brain, outside of the dura. The addition of PEG35 produced similar macroscopic effects, with visible tissue shrinkage observed even at relatively low concentrations. Brains perfused with 5% PEG35 generally appeared more contracted than brains perfused with comparable concentrations of mannitol, and 10% PEG35 produced marked gross shrinkage in several specimens. Taken together, these observations suggest that the addition of both mannitol and PEG35 to the perfusate leads to a dose-dependent tissue shrinkage consistent with dehydration, with PEG35 appearing to produce more visible effects at a lower concentration than mannitol, despite having a lower measured osmolality.

**Figure 3.**
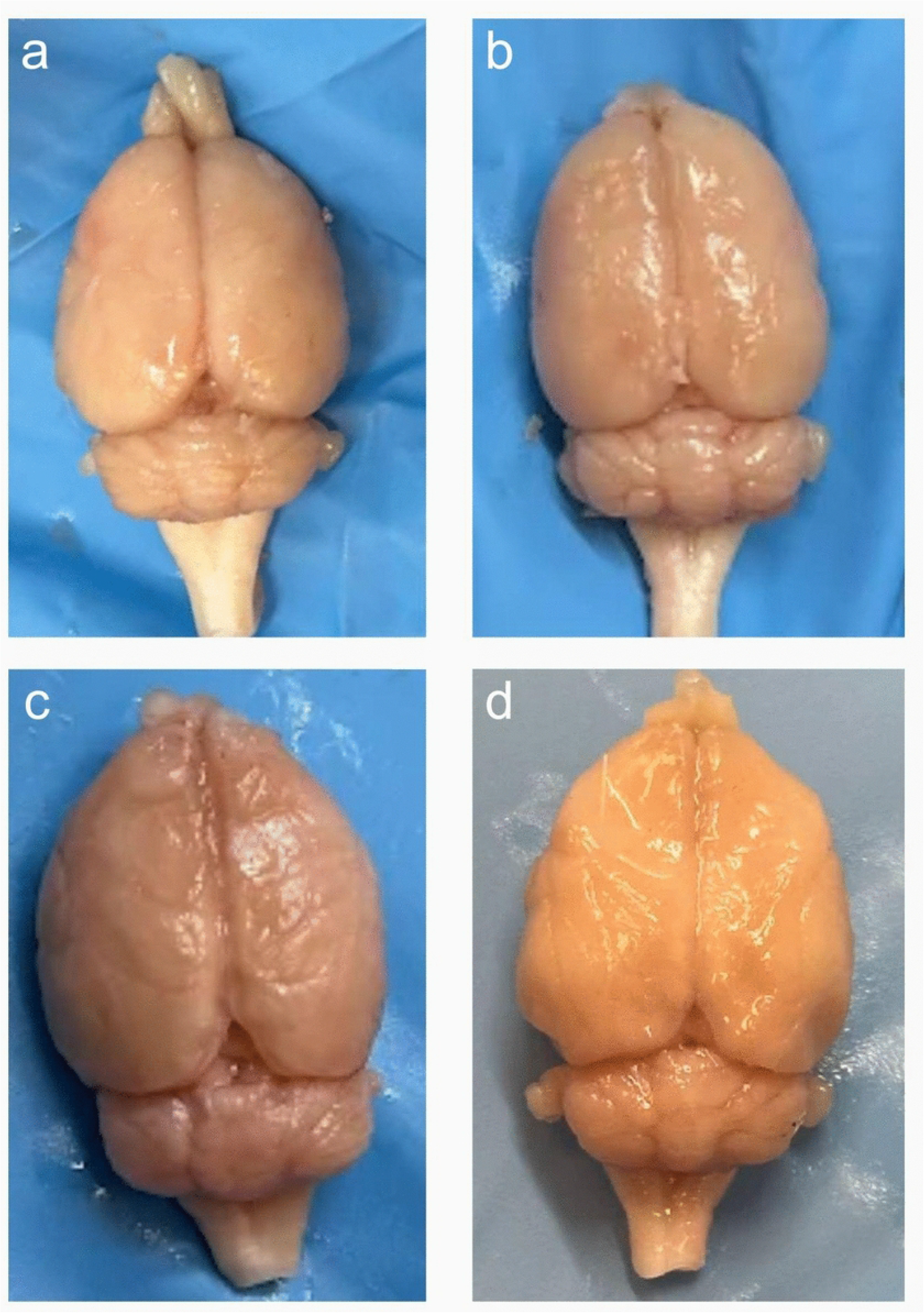
Representative gross brain morphology following perfusion with different fixative formulations. Brains from rats perfused with 20% neutral buffered formalin (NBF) containing 1% glutaraldehyde (GA) (**a**, Rat 11; **b**, Rat 9), 15% mannitol in 20% NBF (**c**, Rat 15), or 10% PEG35 in 20% NBF (**d**, Rat 46). Brains in **c** and **d** show gross tissue shrinkage compared with those in **a** and **b**, consistent with macroscopic dehydration.

### Blood clearance on light microscopy

To determine whether the results on gross examination were due to the actual clearance of intravascular material, we next evaluated the extent of blood removal in LH&E-stained tissue sections. Each WSI received a semi-quantitative vessel clearance grade by two independent reviewers, based on the proportion of vessels free of erythrocytes or other intravascular material (**Figure 4**). These grades had high inter-rater agreement (quadratic-weighted Cohen’s κ = 0.66, n = 56).

**Figure 4.**
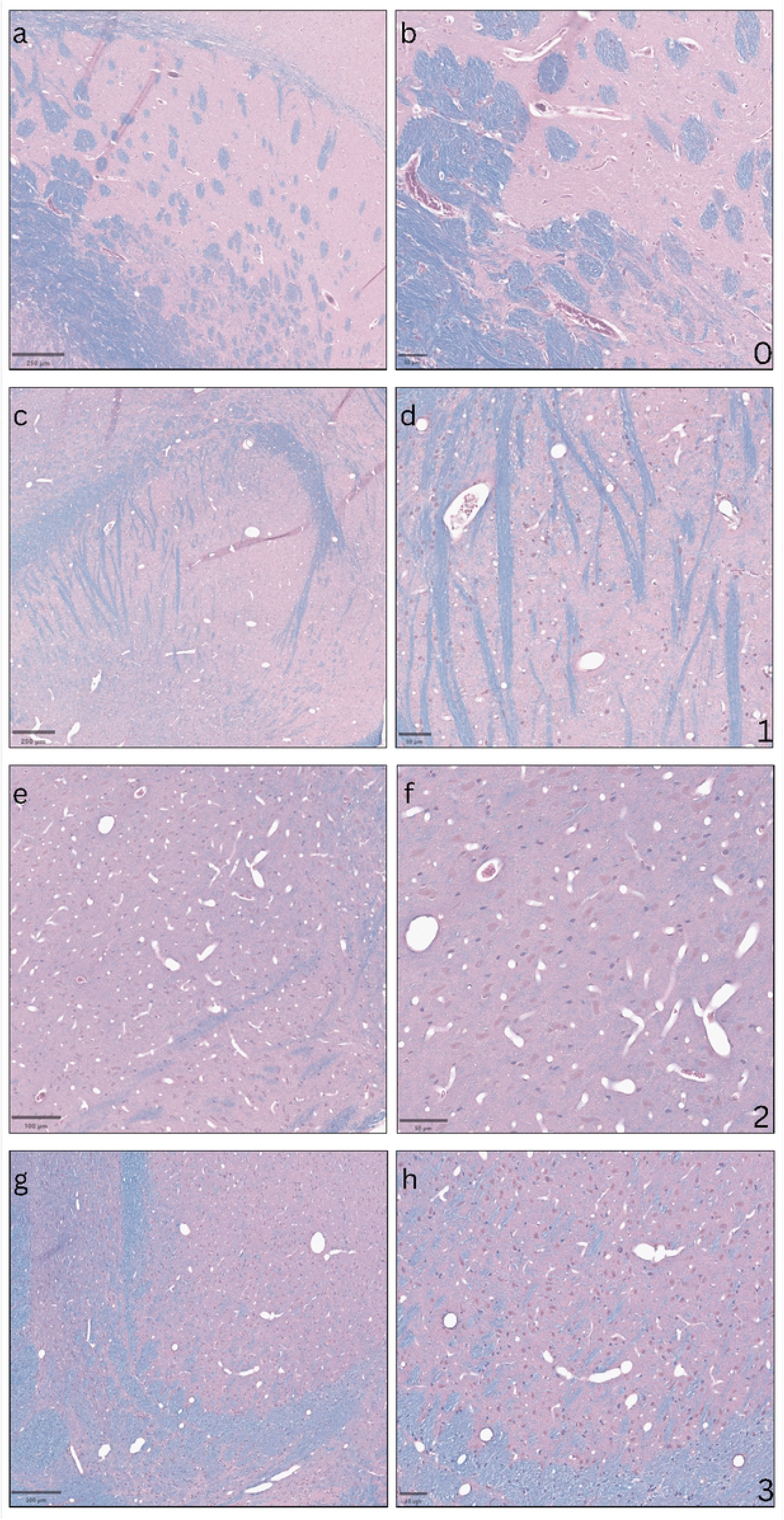
Representative images showing the semi-quantitative grading of vascular blood clearance in whole-slide images. Blood clearance was graded on a 0-3 scale, according to the estimated proportion of vessels cleared of erythrocytes and other intravascular material: 0, <5% clearance (**a**, **b**); 1, 5-50% (**c**, **d**); 2, 50-95% (**e**, **f**); and 3, >95% clearance (**g**, **h**). Panels **a**, **c**, **e**, and **g** show lower-magnification views illustrating the overall distribution of residual intravascular blood, whereas **b**, **d**, **f**, and **h** show corresponding higher-magnification views of blood vessels. Scale bars: 250 µm (**a**, **c**), 200 µm (**g**), 100 µm (**e**), and 50 µm (**b**, **d**, **f**, **h**).

The semi-quantitative grades were next compared with a quantitative measurement of residual RBC burden obtained using a QuPath pixel classifier (**Figure 5** and **Figure 6**). This was calculated in two regions of each WSI, one in the grey matter of the cerebral cortex and one in the mixed white/grey matter of the central region of the brain (**Figure 5**). All of the quantitative measures of residual RBCs that we derived were found to be significantly correlated with the semi-quantitative grades, supporting our use of this ordinal grading scale as a measure of vascular clearance in a given WSI (**Figure 7**).

**Figure 5.**
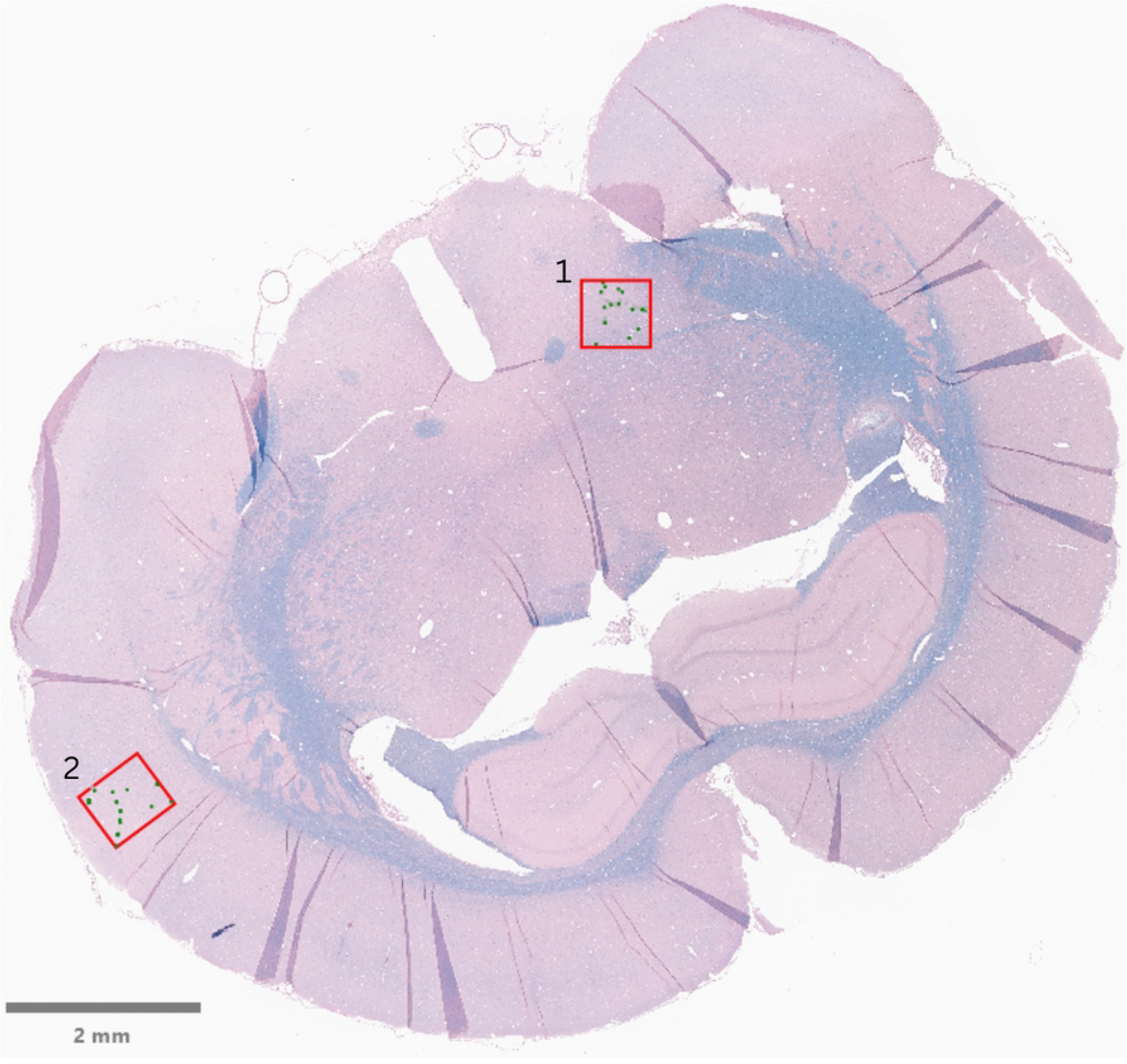
Representative placement of Parent Box 1 and Parent Box 2 in a rat brain histology section (Rat 33). Parent Box 1 was positioned within the central region of the brain sampling an area containing mixed gray and white matter, whereas Parent Box 2 was positioned within the cerebral cortex sampling predominantly gray matter. The red outlines indicate the standardized analysis regions, and green markings represent RBC-positive detections identified by the pixel classifier. Scale bar: 2 mm.

**Figure 6.**
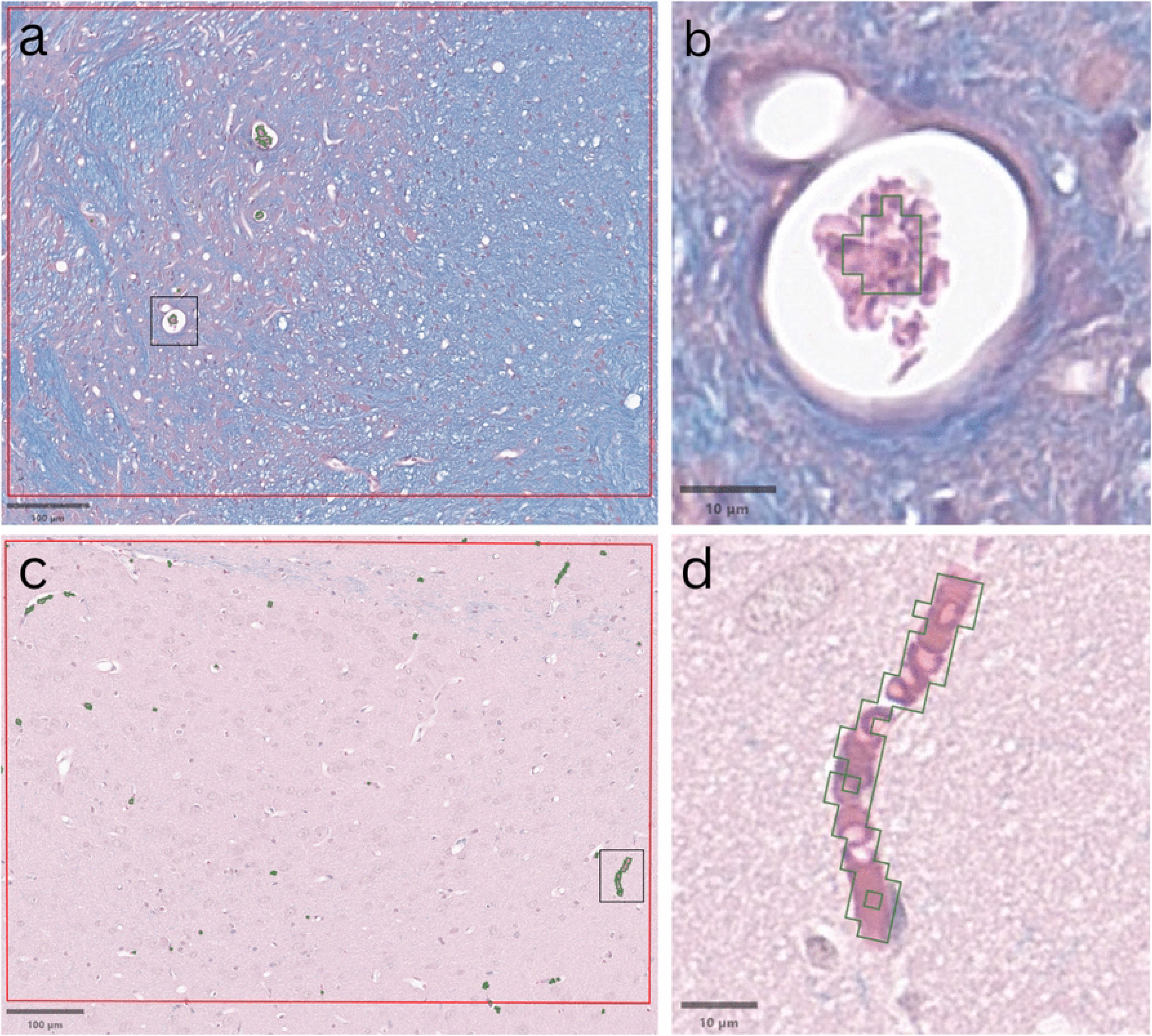
Representative red blood cell detection using the QuPath pixel classifier in Rat 21 (top) and Rat 8 (bottom). (**a**) Representative Parent Box 1 within the central region of the brain, containing mixed gray and white matter; the black box indicates the region shown at higher magnification in **b**. (**c**) Representative Parent Box 2 within the cerebral cortex, containing predominantly gray matter; the black box indicates the region shown at higher magnification in **d**. Dark green regions indicate pixels classified as RBC-positive by the pixel classifier, and red outlines indicate the standardized analysis regions. Scale bars: 100 µm (**a**, **c**) and 10 µm (**b**, **d**).

**Figure 7.**
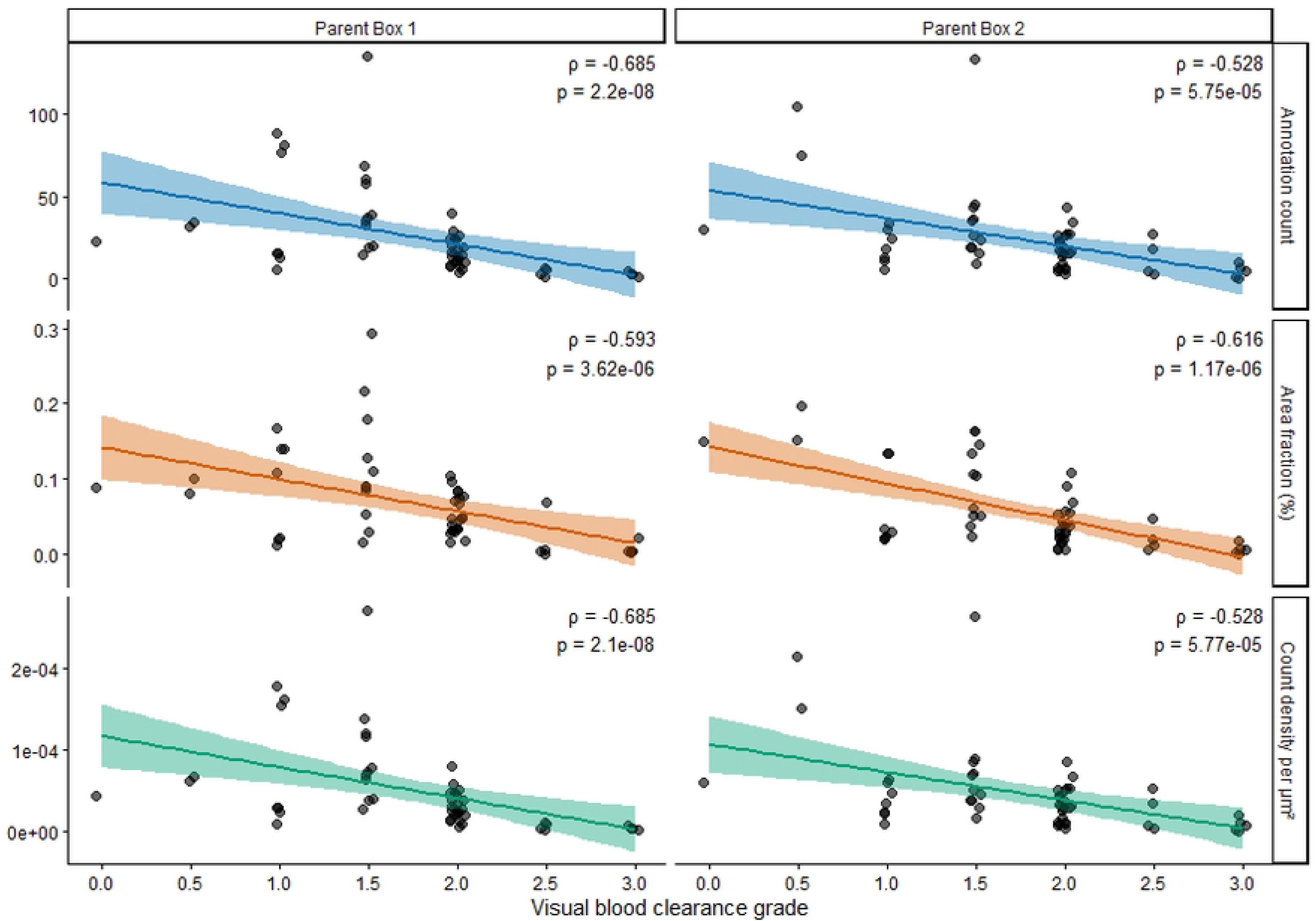
Associations between the semi-quantitative vascular clearance grades and quantitative RBC measurements obtained with the QuPath pixel classifier. The three measurements are shown separately for Parent Box 1 (left column), a central brain region containing both gray and white matter, and Parent Box 2 (right column), a predominantly cortical gray matter region. Rows show the annotation count (i.e., the number of discrete RBC-positive objects detected), RBC-positive area fraction, and the RBC-positive count density. In both of the two types of brain tissue regions analyzed, all three quantitative RBC measures were found to decrease as the semi-quantitative vascular clearance grades increased. Spearman correlation coefficients (ρ) and corresponding p values are shown in each panel. Solid lines indicate fitted linear trends shown for visualization, with shaded bands representing 95% confidence intervals.

Based on these histological observations, we found that brains perfused directly with fixative-containing solutions showed substantial clearance of intravascular blood in many cases. When the rats which received a pre-fixation washout were pooled and compared with those perfused without a washout step, there was no significant difference in the semi-quantitative vessel clearance grades (Mann-Whitney U = 109, p = 0.793; median [IQR]: washout, 1.5 [1.50-2.00]; no washout, 2 [1.50-2.00]). Notably, although residual erythrocytes were present in some specimens across multiple experimental groups, incomplete clearance was not restricted to animals that were perfused with fixative solution alone. These findings indicate that, under the conditions tested, the complete removal of intravascular blood could be effected by the perfusion of fixative alone, and that a preceding PBS washout was not associated with greater vascular clearance.

### Cellular visualization and morphology on light microscopy

We next evaluated whether differences in perfusate composition were associated with differences in cellular visualization and morphology on routine light microscopy. A standardized cortical region of interest was graded by two independent reviewers with two semi-quantitative measures, i.e. (a) the accuracy with which the QuPath cell-detection algorithm outlined cellular profiles (henceforth, shape accuracy) and (b) the visibility of cellular profiles relative to the surrounding tissue (henceforth, cell visibility; **Figure 8**). The agreement between the two reviewers was moderate for shape accuracy (weighted Cohen’s κ = 0.58, n = 56) and substantial for cell visibility (weighted Cohen’s κ = 0.66, n = 56). We found that neither of these measures differed significantly across the full set of treatment groups. The overall Kruskal-Wallis test for shape accuracy was not significant (p = 0.275), and the corresponding test for cellular visibility was also not significant (p = 0.102). Although individual treatment groups varied slightly in their median scores, there was no qualitative pattern indicating superior routine histological preservation quality with a particular conventional fixative or osmotic additive.

**Figure 8.**
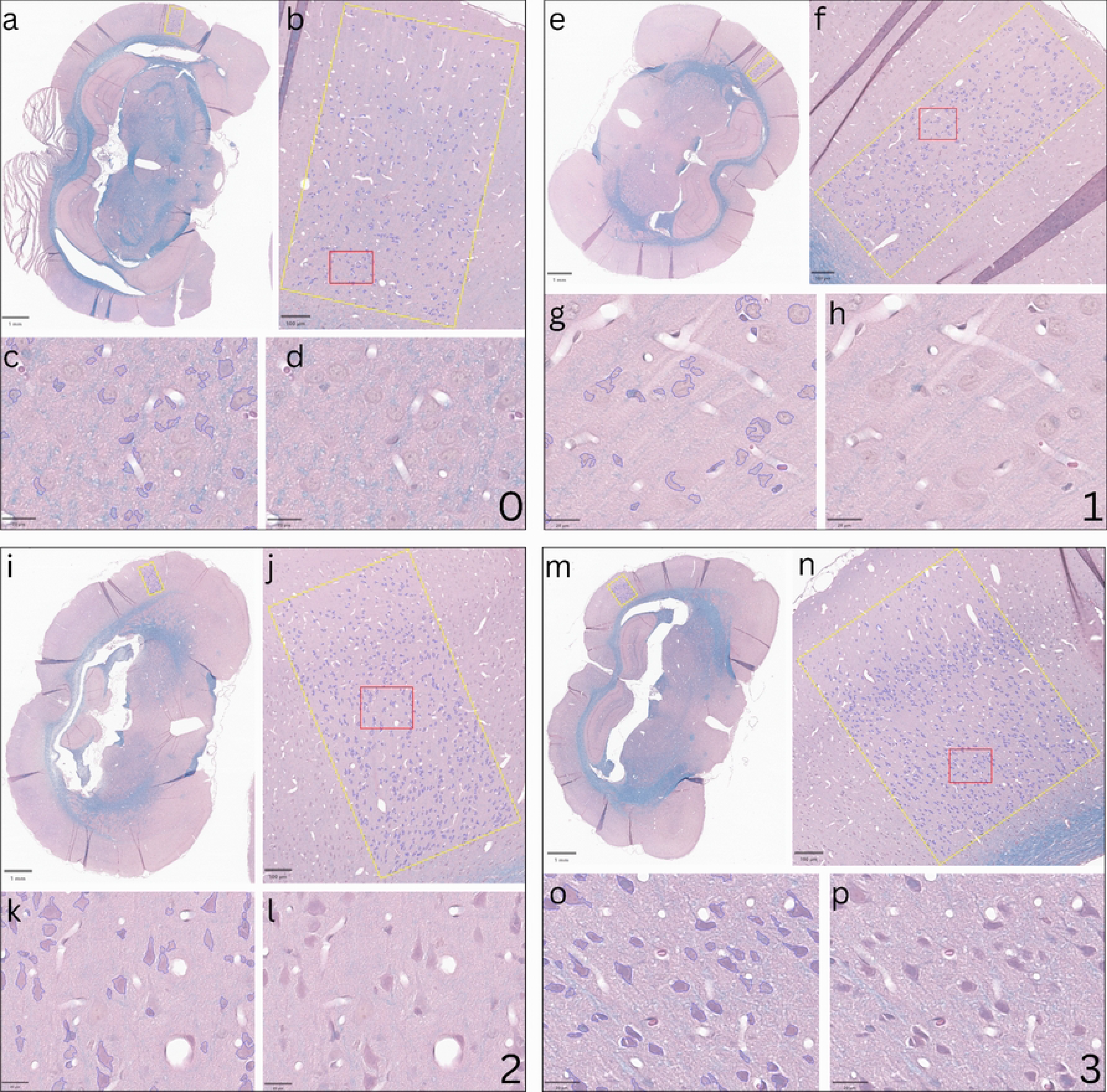
Representative examples of qualitative grading scales showing both the shape accuracy and cell visibility metrics. Although the shape accuracy and cell visibility metrics were graded independently, the two measures were often correlated. In the four representative ROIs shown here, both metrics received the same grade, indicated by the numbers 0-3. The yellow boxes represent a region in the cerebral cortex that is chosen for analysis (**a**, **e**, **i**, **m**). The red boxes indicate where zoomed-in images were taken from for grading (**b**, **f**, **j**, **n**). The tissue within these red boxes is used for the quality grading. Of image pairs **c** and **d**, **g** and **h**, **k** and **l**, and **o** and **p**, the left of the two shows the positive cell detection performed by QuPath, while the right is that portion of the slide without those QuPath annotations. Scale bars: 1 mm (**a**, **e**, **i**, **m**), 100 μm (**b**, **f**, **j**, **n**), 20 μm (**c**, **d**, **g**, **h**, **k**, **l**, **o**, **p**).

We next used QuPath to quantify the shape of detected cellular profiles within the cortical region of interest for each WSI. Detected objects were classified as elongated, round, partially detected, or other (**Figure 9**). The percentage of partially detected objects was significantly associated with both the qualitative shape accuracy and cell visibility ratings (**Figure 10**), supporting its use as a measure of cellular profile delineation.

**Figure 9.**
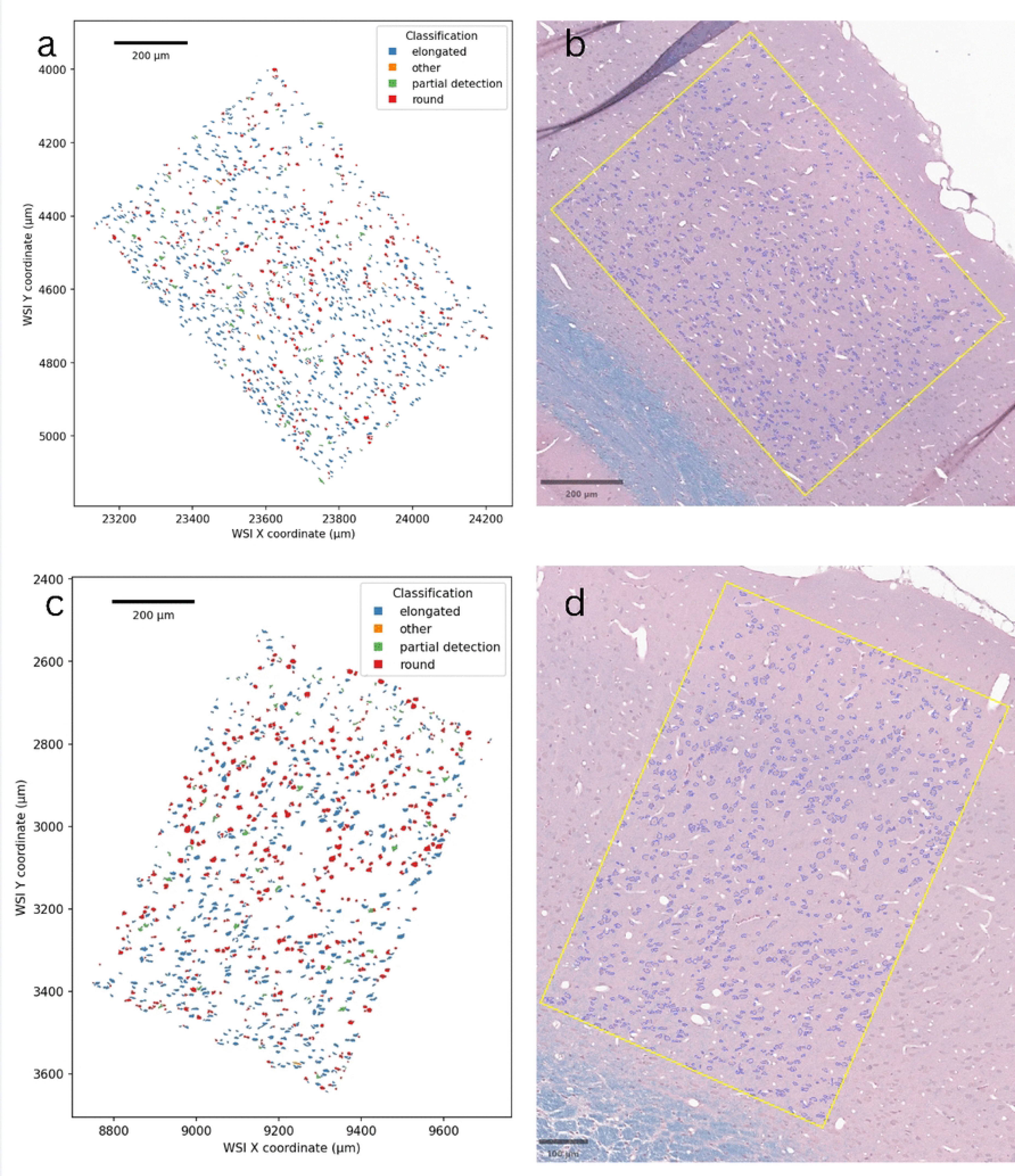
Representative examples of cellular profile shape classification in cortical regions of interest. Images **a** and **c** demonstrate the shape classification of a cortical region of interest. Images **b** and **d** show the site in context with the yellow boundary box and the positive cell detection outlines used to create the figures on the left. Scale bars: 200 μm (**a**, **b**, **c**), 100 μm (**d**).

**Figure 10.**
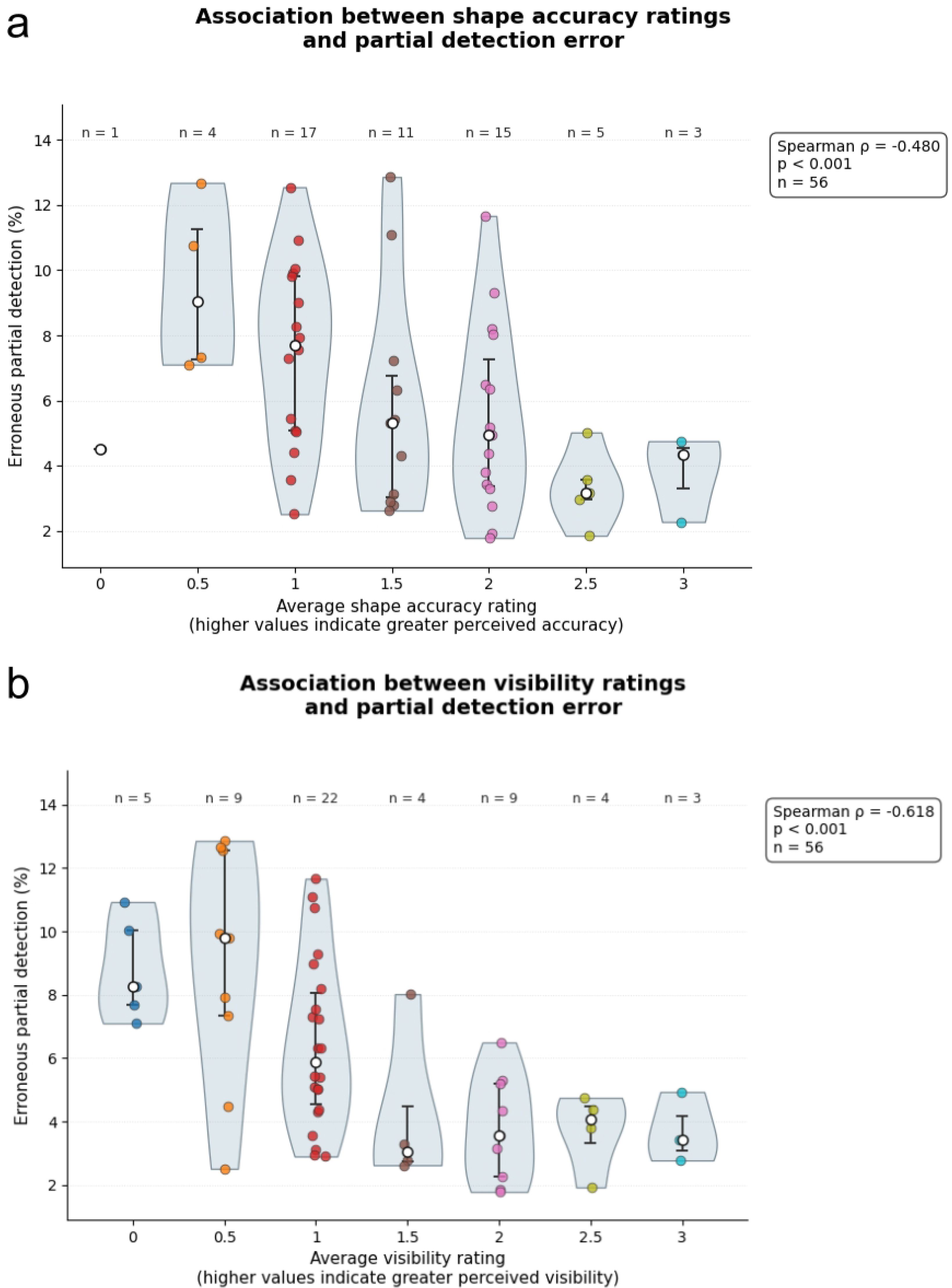
Association between qualitative cellular profile ratings and QuPath-derived partial detection errors. The percentage of QuPath-detected objects classified as erroneous partial detections was compared with the mean reviewer ratings for (a) shape accuracy and (b) cell visibility. Partial detection errors were defined as detected objects with low solidity (<0.75), indicating incomplete or irregular delineation of cellular profiles. Higher reviewer ratings for the quality of the cellular profiles in a WSI were associated with lower percentages of partial detection errors for both shape accuracy (Spearman ρ = −0.480, p < 0.001, n = 56) and cell visibility (Spearman ρ = −0.618, p < 0.001, n = 56). Violin plots show the distribution of partial detection error percentages within each rating category, with individual observations overlaid. The white circles show the median partial detection error percentage per group and the black error bars show the interquartile range.

For the most part, after adjusting for multiple comparison tests, we found no significant relationships between the cellular profile classification metrics and the perfusion parameters, including perfusate volume, perfusion pressure, perfusion rate, perfusion duration, or the concentration of mannitol or PEG35. However, the treatment group was significantly associated with the percentage of partially detected cellular profiles (Kruskal-Wallis test, H(13) = 23.91, p = 0.032, ε² = 0.260). Partial detection was defined as a detected object with low solidity (<0.75), indicating an irregular or incomplete segmentation of the cellular profile. A higher proportion of partially detected objects was interpreted as signifying worse delineation of cellular profiles on the WSI. Dunn’s post hoc pairwise comparisons with a Benjamini-Hochberg correction identified one significant difference among the 91 pairwise comparisons tested. Specifically, the 10% PEG35 in 20% NBF group differed significantly from the 5% mannitol in 20% NBF condition (**Figure 11**). For comparison to a control group, we also show the 20% NBF alone condition, which was qualitatively similar to the 5% mannitol in 20% NBF condition (**Figure 11**).

**Figure 11.**
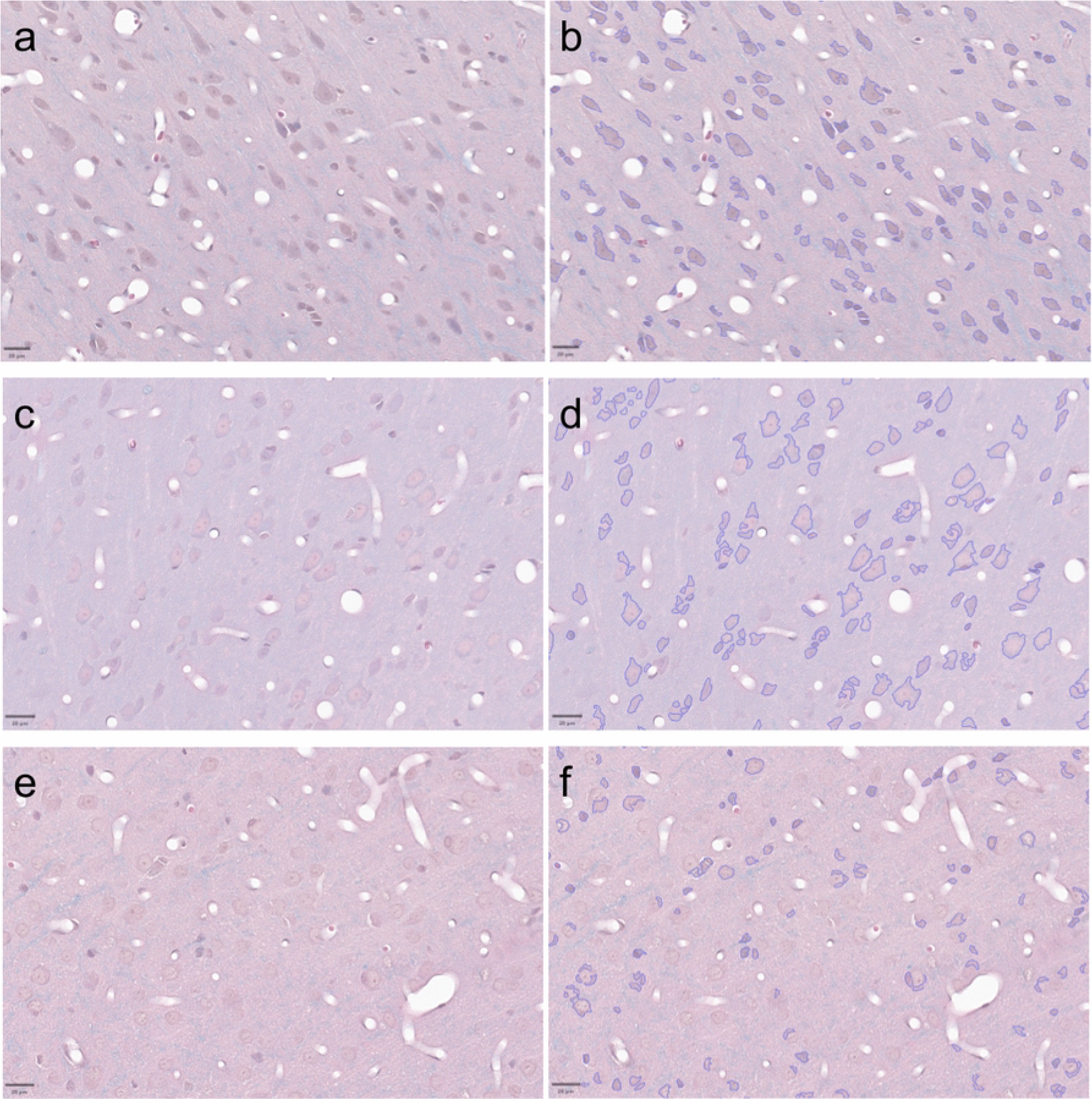
Representative examples of cellular profile detection in selected perfusate conditions. Representative cortical regions are shown from rats perfused with 10% PEG35 in 20% NBF (**a**, **b**), 5% mannitol in 20% NBF (**c**, **d**), or 20% NBF alone (**e**, **f**). Panels **a**, **c**, and **e** show the original LH&E-stained images, whereas **b**, **d**, and **f** show the corresponding QuPath positive-cell detections overlaid on the same regions. Scale bars: 20 µm.

Qualitatively, the WSIs from brains perfused with solutions containing PEG35 appeared to have darker staining, which led to greater contrast of the cells.

### Exploratory ultrastructural assessment

In order to determine whether the broadly similar preservation we observed by light microscopy across different perfusate groups extended to the ultrastructural level, we examined a small subset of specimens by electron microscopy (**Figure 12**). Across the conditions examined, we found that the cellular and neuropil ultrastructure was generally recognizable, with preserved myelinated axons and intracellular organelles visible in representative fields. We did not observe a clear or consistent difference of ultrastructural preservation quality between treatment groups. However, because only one specimen per condition was examined and no blinded scoring was performed, these observations do not permit definitive comparisons between conditions and should be considered exploratory.

**Figure 12.**
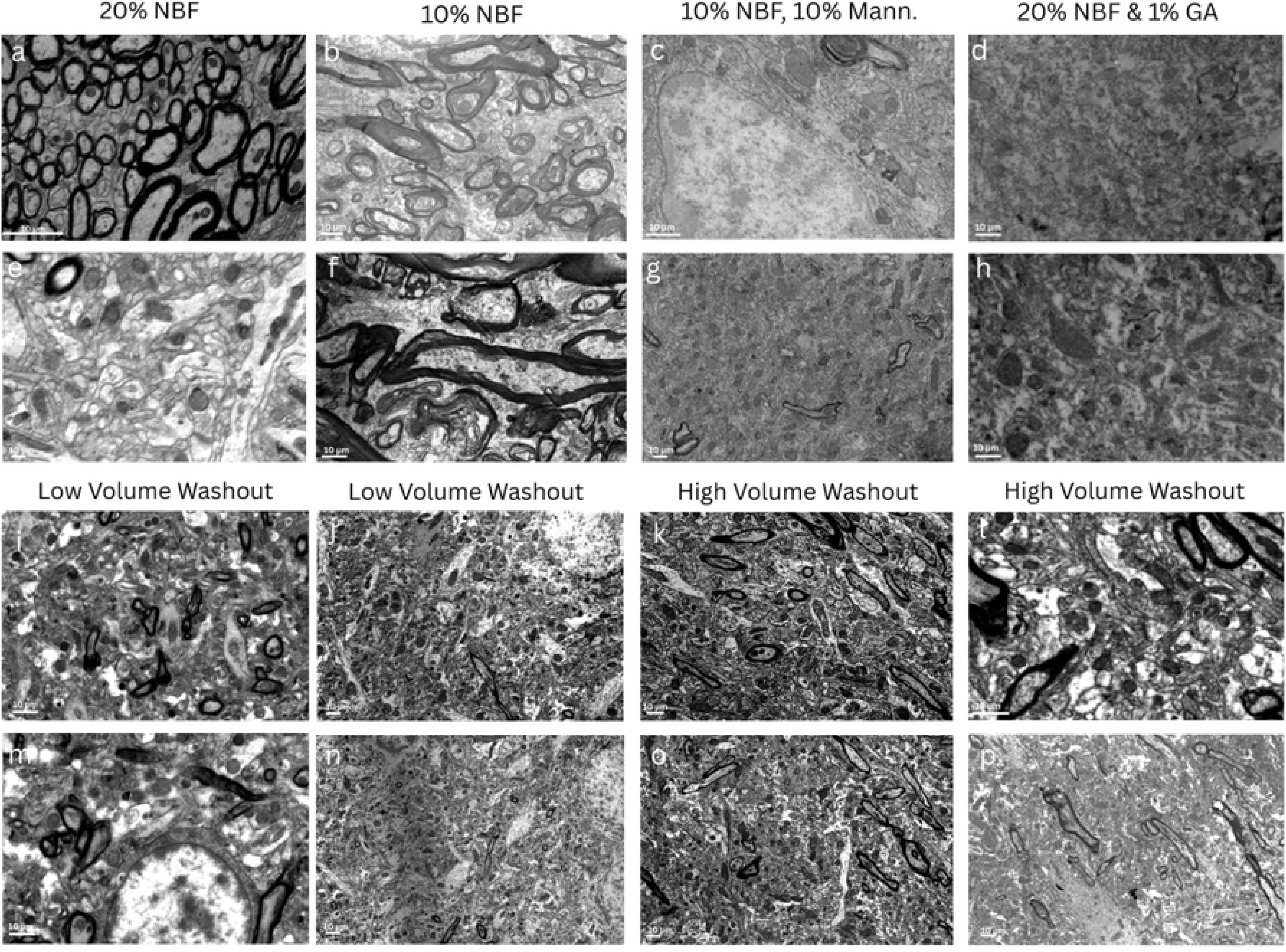
Representative electron micrographs following selected perfusion conditions. Electron micrographs are shown from tissue perfused with 20% NBF (Rat 7; **a**, **e**), 10% NBF (Rat 57; **b**, **f**), 10% NBF with 10% mannitol (Rat 2; **c**, **g**), or 20% NBF with 1% glutaraldehyde (Rat 9; **d**, **h**). The images in the bottom row show preserved tissue from animals receiving either a 50 mL PBS washout (Rat 54; **i**, **j**, Rat 55; **m**, **n**) or a 150 mL PBS washout (**Rat 59**, **k**, **l**, **o**, **p**) before perfusion with 20% NBF. NBF: Neutral buffered formalin, GA: Glutaraldehyde, Mann: Mannitol, PBS: Phosphate buffered saline.

## Discussion

Many of the procedural design choices used in modern perfusion fixation protocols have a long history [1,31]. Parameters that are well-known to play a potential role in mediating preservation quality include the chemical composition, osmotic concentration, and temperature of the fixative, as well as variations in technique such as the duration of ischemia (if any), and the perfusion pressure, flow rate, and duration [32,25,3,33,34]. In this study, we compared several parameters of the chemicals used for transcardial perfusion fixation and quantified their effects on brain preservation quality. Despite substantial variation in the chemical composition of the perfusates used, many of the conditions tested produced broadly similar preservation on routine light microscopy. Brains perfused directly with fixative frequently showed substantial clearance of intravascular blood, and a preceding PBS washout was not associated with detectably greater vascular clearance under the conditions tested. Similarly, we did not identify a clear pattern of differences in histologic preservation quality among the conventional aldehyde fixative formulations examined. Notably, the addition of chemical additives which increased the measured osmolality, i.e. mannitol or PEG35, produced readily apparent tissue shrinkage visible on gross examination. However, these macroscopic changes were not accompanied by similarly large or consistent differences in the routine light microscopy measures examined. Overall, the variations of the perfusate chemical composition we tested were found to have at most a relatively modest effect on tissue quality as measured by histological appearance.

Although we did not identify many differences between treatment groups, we did find a substantial case-to-case variability in perfusion quality, even when the animals were undergoing nominally similar perfusion fixation procedures. Several previous articles have pointed out that technical execution errors are a common reason for poor perfusion and resulting fixation quality, as it is a technically sensitive procedure [1,35–38]. For example, when the cannula is meant to be inserted into the left ventricle, it could instead enter the right ventricle or be inserted at a suboptimal angle, so that it does not point to the aortic outflow tract. Other possible errors include the introduction of air bubbles into the vascular system, using too low of perfusion pressure, or the premature loss of effective cardiac output prior to the initiation of perfusion. In our study, we found that several brains showed signs of inadequate perfusion, such as localized mottling or pinkish-red areas along portions of the lateral cerebral hemispheres. We suspect that this individual case variability was most likely due to differences in the technical execution of each perfusion procedure rather than differences in the composition of the perfusate.

We found that there was comparable blood vessel clearance and routine histological preservation in brains immediately perfused with fixative or brains perfused with a washout solution first. This is consistent with previous reports suggesting that a saline or PBS washout step is not necessary for successful transcardial perfusion fixation in rodents [6,7,10]. We note that there are still some situations in which a washout step could be useful. For example, if the tissue were not to be subsequently perfused with fixative solution, a washout step could be used to remove intravascular material prior to downstream assays such as transcriptomics, where the presence of circulating blood cells could act as a confound. A washout step could also be used to manipulate the blood-brain barrier, osmotic state of the tissue, or extracellular space prior to fixation, which would be expected to lock the tissue into the morphology at the time of fixative perfusion [24]. Outside of applications with a particular need for washout, however, our results did not identify a clear benefit of a routine washout step for the outcomes measured here, while it also increases the complexity of the experiment and the ischemic time prior to preservation.

Our findings are consistent with the broader literature from human brain banking, in which successful perfusion fixation has been reported both with and without a pre-fixation washout step [5]. One possible explanation for the continued widespread use of washout is historical precedent and the benefits of maintaining consistency with previously published studies. Such practices could be described colloquially as “zombie protocols,” -- i.e., methodological steps that persist largely through historical precedent despite limited direct evidence that they are *sensu stricto* necessary.

The addition of both of the osmotically active agents to the fixative perfusate that we tested, mannitol and PEG35, produced dose-dependent brain tissue shrinkage. The more pronounced gross tissue shrinkage that we observed with solutions containing PEG35, despite these having a lower measured osmolality than the solutions containing mannitol, may be due to the strong colloid osmotic effects of PEG35, although the mechanism was not examined in this study. Previous research has found that changing the osmolality of the fixative perfusate and adding a colloid can have significant effects on cellular volume, extracellular space, and tissue ultrastructure [23,28]. We included these additives to model one approach to human postmortem brain perfusion, where osmotic agents and colloids have been proposed to ameliorate the ischemia-associated vasogenic edema and other forms of perfusion impairment [22,26]. Notably, we recently found that human and canine brains perfused with fixative solution containing mannitol and PEG35 could show relatively favorable measures of perfusion quality, while also exhibiting substantial ultrastructural artifacts, attributed to osmotic shock [39]. Here, we found that osmotic additives generally produced much more pronounced effects on gross tissue morphology than on the routine light microscopy measures examined. One exception was the 10% PEG35 group, which showed a lower proportion of partially detected cellular profiles on light microscopy than one other group. Notably, specimens perfused with PEG35 also appeared qualitatively darker and showed greater contrast between cellular profiles and the surrounding neuropil. This could be due to dehydration or due to changes in the appearance of the surrounding neuropil, but the basis of this staining difference was not assessed in our study. Together, our findings suggest that osmotic additives may be useful when a particular tissue geometry or vascular effect is desired, but that they must be used judiciously, if at all, because these agents can potentially lead to morphological artifacts at higher concentrations.

## Conclusions

Across the perfusate formulations we examined, substantial differences in chemical composition generally produced only modest differences in preservation quality as visualized by routine light microscopy. A pre-fixation PBS washout was not found to be necessary to achieve substantial vascular blood clearance under the conditions we tested. The addition of mannitol or PEG35 produced gross tissue shrinkage without substantial corresponding alterations in histological preservation, with the exception of a lower proportion of partially detected cellular profiles in the 10% PEG35 group compared with one other condition. The substantial variability in perfusion quality observed among animals undergoing nominally similar procedures suggests that technical execution is also a critical determinant of brain preservation. Taken together, our findings support the importance of empirically evaluating different possible steps in a perfusion fixation protocol, as theoretical considerations alone may be insufficient to determine which protocol modifications meaningfully affect preservation quality.

## Abbreviations

BBB: blood-brain barrier
EM: electron microscopy
GA: glutaraldehyde
LH&E: Luxol fast blue, hematoxylin, and eosin
MW: molecular weight
NBF: neutral buffered formalin
PBS: phosphate-buffered saline
PEG35: polyethylene glycol, 35 kDa
PFA: paraformaldehyde
RBC: red blood cell
ROI: region of interest
WSI: whole-slide image.

## Author contributions

Autumn Beck: Investigation, Formal analysis, Writing - original draft, Writing - review & editing. Andria Slaughter: Investigation, Formal analysis, Writing - original draft, Writing - review & editing. Sarah Sedgewick: Formal analysis, Writing - original draft, Writing - review & editing. Macy Garrood: Formal analysis, Writing - review & editing. Katelyn Hedden: Formal analysis, Writing - review & editing. Andrew T. McKenzie: Conceptualization, Methodology, Formal analysis, Supervision, Project administration, Writing - original draft, Writing - review & editing.

## Acknowledgements

We would like to acknowledge the Neuropathology Brain Bank & Research CoRE at the Icahn School of Medicine at Mount Sinai for their histology and tissue processing services. The Icahn School of Medicine at Mount Sinai provided access to library resources.

## Data availability

Whole slide image data can be accessed in a public repository on Zenodo, available here: https://doi.org/10.5281/zenodo.22210994 and here: https://doi.org/10.5281/zenodo.22217323. Electron microscopy data can be accessed in a public repository on Zenodo, available here: https://doi.org/10.5281/zenodo.22239137. Rating data and statistical analysis can be accessed here: https://zenodo.org/records/22775542. Code used for data analysis and a spreadsheet version of the rating data and analysis workbook is available here: https://github.com/andymckenzie/comparative-rodent-histology. The QuPath RBC detection model is available at this DOI: https://zenodo.org/records/22237547. The QuPath positive cell detection code is available at this DOI: https://doi.org/10.5281/zenodo.22236780.

